# Patterns of temporal and enemy niche use by a community of leaf cone moths (*Caloptilia*) coexisting on maples (*Acer*) as revealed by metabarcoding

**DOI:** 10.1101/094417

**Authors:** Ryosuke Nakadai, Atsushi Kawakita

## Abstract

The diversity of herbivorous insects is often considered a function of host plant diversity. However, recent research has uncovered many examples of closely related herbivores using the same host plant(s), suggesting that partitioning of host plants is not the only mechanism generating diversity. Herbivores sharing hosts may utilize different parts of the same plant, but such resource partitioning is often not apparent; hence, the factors that allow closely related herbivores to coexist are still largely undetermined. We examined whether partitioning of phenology or natural enemies may explain the coexistence of leaf cone moths (*Caloptilia*; Gracillariidae) associated with maples *(Acer;* Sapindaceae). Larval activity of 10 sympatric *Caloptilia* species found on nine maple species was monitored every 2–3 weeks for a total of 13 sampling events, and an exhaustive search for internal parasitoid wasps was conducted using high-throughput sequencing. Blocking primers were used to facilitate the detection of wasp larvae inside moth tissue. We found considerable phenological overlap among *Caloptilia* species, with two clear peaks in July and September–October. Coexisting *Caloptilia* species also had largely overlapping parasitoid communities; a total of 13 wasp species belonging to four families attacked *Caloptilia* in a non-specific fashion at an overall parasitism rate of 46.4%. Although coexistence may be facilitated by factors not accounted for in this study, it appears that niche partitioning is not necessary for closely related herbivores to stably coexist on shared hosts. Co-occurrence without resource partitioning may provide an additional axis along which herbivorous insects attain increased species richness.

## Introduction

Host plant diversity is arguably the primary factor that drives the diversity of herbivorous insects on earth (Novotny *et al.* 2006). Because herbivore species are usually specialized to a narrow taxonomic group of plants, ecological speciation as the result of a shift to a new host (host-shift-driven speciation) is often considered a major driver of herbivorous insect diversification (Feder *et al.* 1988; Hawthorne & Via 2001; Nosil *et al.* 2002; Malausa *et al.* 2005). For example, a classic study by Farrell (1998) showed that among the Phytophaga beetles, lineages that use angiosperms as hosts are more species rich than are those that use gymnosperms, suggesting that the diversity of angiosperms has facilitated host-shift-driven diversification of the beetles that feed on them. However, different views about the effects of host plants in generating herbivorous insect diversity have arisen (Rabosky 2009; Kisel *et al.* 2011; Nyman *et al.*2012) as studies increasingly document examples where closely related herbivores share the same host plants (Nyman *et al.* 2010; Imada *et al.* 2011; Nakadai & Kawakita 2016). One hypothesis is that host plants facilitate the coexistence of species that have already diverged, and a shift to a new host is not necessary at the time of speciation(Rabosky 2009; Nakadai & Kawakita 2016); however, the exact role of host plants in facilitating local coexistence has not been well studied. Studying the mechanisms that permit local coexistence of closely related herbivores is important, as both the number of locally coexisting species and the mean geographical range size are significant estimators of global species diversity (Storch *et al.* 2012)

Correlation between host plant diversity and herbivorous insect diversity is often confirmed at the local community level (Siemann *et al.* 1998; Borer *et al.* 2012). This indicates that the use of different host plants is important for niche partitioning and species coexistence (MacArthur & Levins 1967; Benson 1978). However, there are many examples where closely related herbivores overlap in their use of host plants. Herbivores that share hosts sometimes partition resources by using different parts of the same plant or leaves of different ages (Benson 1978; Bailey *et al.* 2009; Condon *et al.* 2014). For example, Benson (1978) confirmed niche partitioning among *Heliconiini* butterflies along three different niche axes (plant species, plant habitat, and plant part). However, in many instances, closely related herbivores co-occur on the same host without any apparent means of resource partitioning (Strong *et al.* 1982), indicating that there are other factors besides resource partitioning that facilitate coexistence of species sharing the same host plant.

One mechanism that allows coexistence of species with similar resource use is phenological partitioning. For example,the geometrid winter moth *Inurois punctigera* has two allochronic races that coexist stably without partitioning resources; allochrony is even postulated as the direct cause of divergence in this case (Yamamoto & Sota 2009). Alternatively, species that share the same food resources can have different natural enemies and thereby occupy non-overlapping niches. Condon *et al.* (2014) demonstrated that species-specific parasitoids increase the niche diversity of *Blepharoneura* flies co-existing on the same-sex flowers of curcurbit host plants. Also, the more than 20 *Andricus* gall wasp species that coexist on shared oak hosts display remarkable diversity of gall forms; because gall morphology is a major determinant of parasitoid community structure, differences in natural enemies also provide a comprehensive explanation for the coexistence of multiple gall wasp species on oaks (Bailey *et al.* 2009). However, analysis of parasitoid communities among closely related herbivores is still limited, and our understanding of the role of natural enemies will increase with additional data.

In this study, we examined whether differences in phenology or natural enemies explain the coexistence of closely related herbivorous insects on shared host plants. We focused on interactions between a group of leaf cone moths (*Caloptilia*, Gracillariidae) and their maple hosts (*Acer*, Sapindaceae) because previous studies have identified multiple pairs of species that occur sympartrically with a great deal of overlap in host use (Kumata 1982; Nakadai & Murakami 2015). With 124 species, the genus *Acer* is one of the most species-rich groups of trees in the northern hemisphere, particularly in the temperate regions of East Asia, eastern North America, and Europe (van Gelderen *et al.* 1994). In temperate Japan, as many as 20 *Acer* species can occur in a single location (Nakadai et al. 2014), which may host to up to 10 sympatric *Caloptilia* species, as predicted from the geographic distribution of leaf cone moths (Nakadai & Kawakita 2016). Twenty-eight *Acer* species occur in Japan (Nakadai et al. 2014), and a previous study confirmed 14 Acer-feeding *Caloptilia* species; 13 of the 14 *Caloptilia* species formed a monophyletic group, together with a *Toxicodendron-feeding Caloptilia*, in the global *Caloptilia* phylogeny and thus are very closely related (Nakadai & Kawakita 2016). We investigated the phenology (i.e., temporal niche) and parasitoid community (i.e., enemy niche) of locally co-occurring, maple-feeding *Caloptilia* species by sampling *Acer* leaves containing *Caloptilia* larvae every 2-3 weeks for a total of 13 sampling events, yielding 274 moth larvae. Species identification of moth larvae and detection of internal parasitoids were based on a simultaneous barcoding (metabarcoding) approach using high-throughput sequencing with the aid of *Caloptilia*-specific blocking primers that effectively reduced the number of redundant moth reads.

## Materials and Methods

### Study materials

The genus *Caloptilia* is globally distributed and includes nearly 300 described species, of which 27 feed on maples (De Prins & De Prins 2015; Kawahara *et al.* 2016). In Japan, there are 51 described *Caloptilia* species feeding on 21 host plant families (Kumata *et al.*2013). Eleven of these species are known to use *Acer*, which is the most common host plant genus for Japanese *Caloptilia* (Fig. 1) (Kumata *et al.* 2013). Three additional *Caloptilia* species were newly found feeding on *Acer* in recent years. Most of the Japanese *Caloptilia* moths are multivoltine (Kumata *et al.* 2013). The feeding habits of the larvae change dramatically between the early and late developmental stages. Upon hatching, larvae mine the surface layer of the leaf, until the third instar. They then exit the mine and roll the edge of the leaf to form a cone, within which they feed externally until the final instar (Kumata *et al.* 2013). Some species are leaf-gallers or blotch-miners at the final instar and do not roll leaves (Nakadai & Kawakita 2016). Previous phylogenetic analysis of *Caloptilia* moths showed that the Japanese species of *Caloptilia* moths that feed on maples are closely related (Fig. S1) (Nakadai & Kawakita 2016).

**Figure 1.**
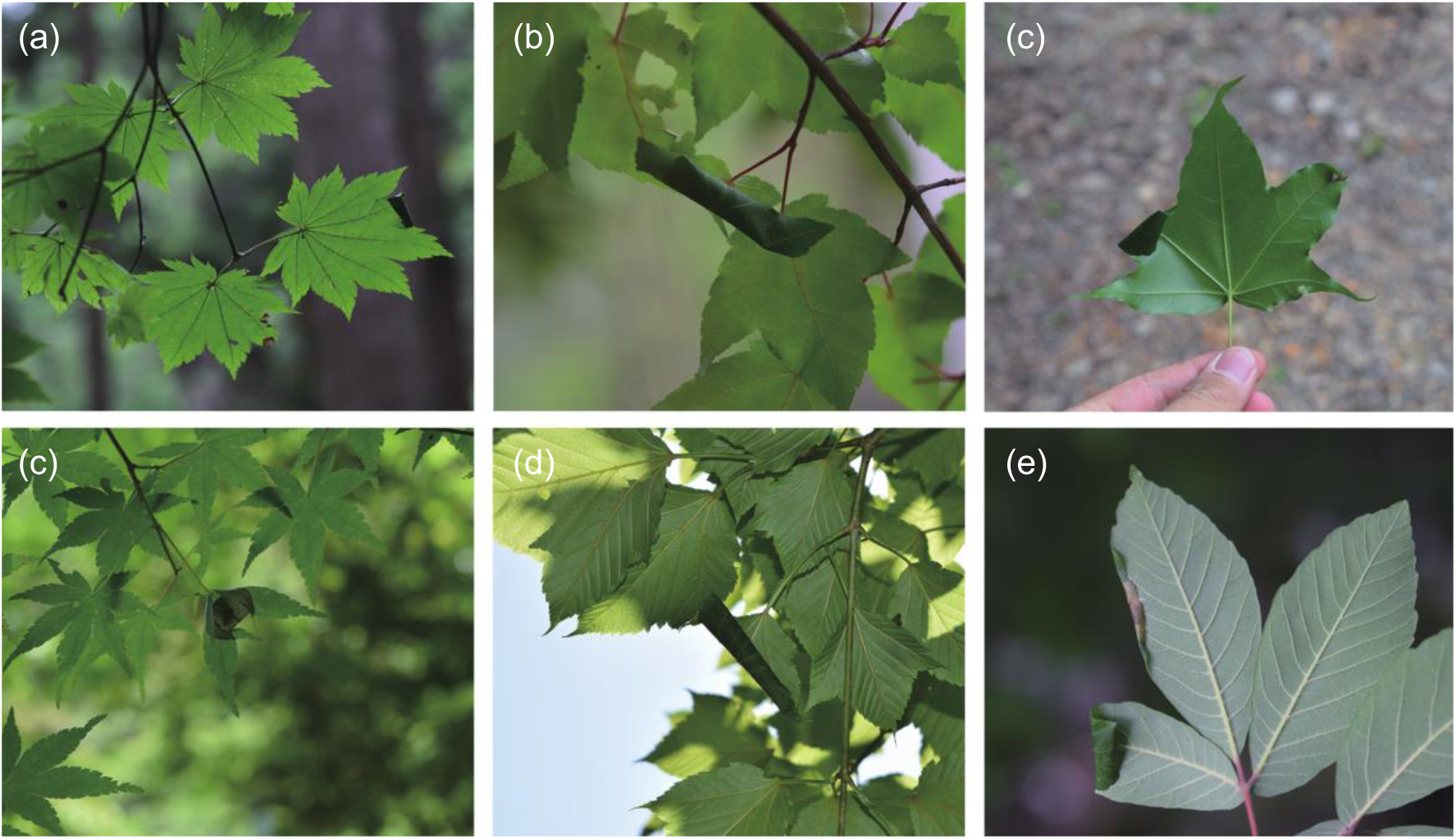
Leaves rolled by leaf cone moths on Japanese maples; moth species are very difficult to identify based solely on the morphology of rolled leaves and larvae. (a) *Acer japonicum*, (b) *A. crataegifolium*, (c) *A. pictum*, (d) *A. palmatum*, (e) *A. rufinerve*, (f) *A. maximowiczianum*.

### Study sites

We conducted field surveys in a natural temperate forest at Ashiu Forest Research Station of Kyoto University (35°18′ N, 135°43′ E). The forest is dominated by *Fagus crenata* and *Quercus crispula* above 600 m elevation and Q. *serrata, Q. salicina,* and *Ilex pedunculosa* below 600 m (Ashiu Forest Research Station 2015). The average annual temperature for 1981–2010 was 12.1°C, and the average annual rainfall for 1981–2010 was 2,257 mm (Ashiu Forest Research Station 2015).

### Acer species

Fourteen *Acer* species have been confirmed in the Ashiu forest (Yasuda & Nagamasu 1995), but because four of these species are rare (*A. cissifolium, A. diabolicum, A. palmatum,* and *A. tenuifolium)*, we targeted the following ten species: *Acer amoenum, A. carpinifolium*, *A. crataegifolium*, *A. japonicum*, *A. maximowiczianum*, *A. micranthum*, *A. nipponicum* subsp. *nipponicum, A. pictum, A. rufinerve,* and A. *sieboldianum*.

### Sampling and species identification of leaf cone moths and the search for internal parasitoid wasps

*Caloptilia* moths feeding on *Acer* trees were sampled every 2–3 weeks by searching for active larvae in leaf rolls (i.e., fourth or fifth instar) on the foliage of 10 *Acer* species from mid-May to mid-November of 2015 (Fig. 3). We sampled only larvae in leaf rolls because some leaf-mining larvae die early due to inconsistency between maternal oviposition and larval performance, and host use cannot be assessed precisely in such cases. This also enabled us to avoid sampling artifacts caused by the difficulty of conducting an exhaustive search for leaf miners. To standardize sampling effort, we sampled *Caloptilia* moths from branches with a diameter of 2.1 ± 4 mm from five individuals of each tree species. After sampling, moths were preserved in 99.5% ethanol and stored at −20°C.

Delimitation of species was based on sequences of the mitochondrial cytochrome oxidase subunit I (COI) gene for all samples. We extracted genomic DNA using the NucleoSpin Tissue Kit (Macherey-Nagel, Düren, Germany). To simultaneously perform species identification of larvae and an exhaustive search for internal parasitoid wasps by high-throughput sequencing, we amplified the mitochondrial cytochrome oxidase I gene using the primers mlCOIlintF (Leray *et al.* 2013),5′-GGWACWGGWTGAACWGTWTAYCCYCC-3′, and HCO2198 (Folmer *et al.*1994), 5′-TAAACTTCAGGGTGACCAAAAAATCA-3′, which produced fragments with a standard sequence length of 313 base pairs. The COI region has been adopted as the standard ‘taxon barcode’ for most animal groups (Hebert *et al.* 2003) and is by far the most represented in public reference libraries. This primer set has performed well in previous studies that exhaustively searched for animal phyla (Leray *et al.* 2013; Brandon-Mong *et al.* 2015). We employed a two-step tailed PCR approach to conduct massively parallel paired-end sequencing (2 × 250 bp) on the MiSeq platform (Illumina, San Diego, CA, USA) (FASMAC Co., Ltd., Kanagawa, Japan).

The first PCR was carried out in a total volume of 10 μl including 0.5 ng of DNA, 5 μl of Kapa HiFi Hotstart ReadyMix (Kapa Biosystems, Wilmington, MA, USA), and 0.3 μM each of forward and reverse primers. We also added the blocking primer (Vestheim & Jarman 2008) for *Caloptilia* moths at eight times the concentration of versatile primers (see the next section for details about the blocking primer). The protocol for the first PCR was 2 min at 95°C, followed by 35 cycles of 20 s at 98°C, 15 s at 67°C, 30 s at 52°C, and 30 s at 72°C, with a final extension at 72°C for 1 min. Purification of the first PCR products was done with Agencourt AMPure XP (Beckman Coulter, Brea, CA, USA). The second PCR was then carried out in a total volume of 10μl including 1 μl of the template DNA amplified in the first PCR, 5 μl of Kapa HiFi Hotstart ReadyMix, and 0.3 μM each of forward and reverse primers for the second PCR. The protocol for the second PCR was 2 min at 95°C, followed by 12 cycles of 20 s at 98°C, 30 s at 60°C, and 30 s at 72°C, with a final extension at 72°C for 1 min. Purification of products of the second PCR was also done with Agencourt AMPure XP. PCR products were normalized and pooled. We normalized PCR products after quantifying them with a Nano Drop ND-1000 (Thermo Fisher Scientific, Waltham, MA, USA).

### Design of Caloptilia moth-specific annealing blocking primer

The bodies of *Caloptilia* moths include both their own tissues and, occasionally, those of internal parasitoids; the ratio of parasitoid tissue to moth tissue is very small. Thus, the amplification of *Caloptilia* moth COI sequences must be suppressed to allow detection of the sequences of internal parasitoids. We used the blocking primer approach for this purpose (Vestheim & Jarman 2008). A blocking primer is a modified primer that overlaps with one of the binding sites of the versatile primer. Blocking primers are usually designed for only one species (Leray *et al.* 2015), so it is difficult to apply them to multiple closely related species, as in this study, where we were unable to identify the larvae morphologically. We designed the blocking primer 5′-CCCCCCHCTTTCATCWAAYATYGCHCATRGWGGWAGATC-3′ to block sequences of *Caloptilia* moths feeding on *Acer* based on known information about the COI sequences of the moths and already-confirmed parasitoids (Table S4). The blocking primer overlaps six bases with mlCOIlintF. The blocking primer was modified at the 3′-end with a Spacer C3 CPG (three hydrocarbons) to prevent elongation without affecting its annealing properties (Vestheim & Jarman 2008). The performance of the blocking primer for *Caloptilia* moths feeding on *Acer* was also tested (Supplemental files 2).

### Analysis of sequencing data

We extracted the reads, which fully contain both primer sequences, from the output FastaQ files of Miseq using FastX Toolkit (ver. 0.0.13.2; Gordon & Hannon 2010). The remaining adapter, primer region, and the last nucleotide were trimmed from the reads using FastX Toolkit. Additionally, the reads were quality filtered using Sickle (ver. 1.33; Joshi & Fass 2011) with a minimum Sanger quality of 20 and a minimum length of 100. Paired reads were assembled using FLASH (ver. 1.2.10; Magoč & Salzberg 2011) with a minimum overlap of 20, and then transformed into Fasta format using FastX Toolkit. A de novo chimera removal was performed using the UCHIME algorithm (Edgar *et al.* 2011) of USEARCH (ver. 8.0.1623_i86linux64; Edgar 2010). Duplicate and singleton reads were removed using USEARCH. We used the UPARSE-OTU algorithm (Edgar 2013) of USEARCH for clustering OTUs with an identity threshold of 97%. Thereafter, taxonomic assignment of individual OTUs was performed by BLAST+ (ver. 2.2.29; Camacho *et al.* 2009).

Subsequently, non-target OTUs (organisms other than *Caloptilia* moths and Hymenoptera) were removed, and OTUs were filtered with a sequence length of 313 ± 15 base pairs. We found some artifacts in the region overlapping with the blocking primer, so after excluding that region, we clustered OTUs manually with an identity threshold of 97% using MEGA (ver. 6.06; Tamura *et al.* 2013) once again. Additionally, rare OTUs, whose total read number in the whole sample was under 100, were also removed. The most abundant OTU of *Caloptilia* moth in each sample was used for the identification of *Caloptilia* moths, and the OTUs of internal parasitoid wasp over 10 reads in each sample were defined as states of presence.

### Statistical analysis

To assess niche use trends (i.e., niche partitioning or overlap), we calculated the degree of niche overlap using the Pianka (Pianka 1973) and Czekanowski (Feinsinger *et al.* 1981) indices, which are common measures of niche overlap, and compared the observed values with the expectations of null models. Null model-based analyses are one of the most general approaches for assessing niche use trends (Gotelli 2001; Albrecht & Gotelli 2001). As a well-known example, Lawlor (1980) tested the patterns of niche use among 10 North American lizards using four types of null model (the algorithms RA1–4) and confirmed significantly low overlap in resource use, suggesting that interspecific competition plays an important role in constructing community structure. We used the R package EcoSimR (Gotelli *et al.* 2013) for the enemy niche and the program TimeOverlap, which is based on the algorithm ROSARIO (Castro-Arellano *et al.* 2010), for the temporal niche. Because of the sequential and continuous nature of time, a different kind of randomization model is required for the temporal niche (Castro-Arellano *et al.* 2010). Samples that did not host parasitoid wasps were excluded from the analysis of enemy niche. In all tests, the two-tailed probability of the observed value was calculated based on 10,000 randomizations. We employed Lawlor’s (1980) algorithm RA3 for constructing null models. Additionally, we calculated the standardized effect size (SES) as the observed test statistic minus the mean of the null distribution, divided by the standard deviation of the null distribution (Nakadai &awakita 2016). This null model approach is commonly used for expressing biological differences regardless of the units of the indices (McCabe *et al.* 2012). Moreover, to reveal the relationships among the niches of *Caloptilia* moths, we also tested the correlations between overlaps of three niches (resource, temporal, and enemy) and phylogenetic distances between *Caloptilia* moths using Mantel tests. Phylogenetic distances among *Caloptilia* moths were calculated from the phylogeny of Nakadai and Kawakita (2016).

## Results

A total of 274 *Caloptilia* larvae were sampled from nine *Acer* species (all target species except *A. carpinifolium*) in 13 seasonal sampling events. Through high-throughput sequencing, we obtained 5,423,301 reads and 152 OTUs after bioinformatics preprocessing, and 10 OTUs of *Caloptilia* moths and 13 OTUs of internal parasitoid wasps after manual filtering (Table S1). The OTUs include 10 *Caloptilia* species (*C. acericola*, *C. aceris*, *C. gloriosa*, *C. heringi*, *C. hidakensis*, *C. monticola*, *C. semifasciella*, *C*. sp. 1, and *C*. sp. 3) and 13 internal parasitoid wasps (Braconidae, Eulophidae, Icheumonidae, and Trichogrammatidae) (Figs. 3, 4, and Table S1). The names of *Caloptilia* moths were matched with those found by Nakadai and Kawakita (2016). Each *Caloptilia* species uses 1–3 *Acer* species, with an average of 1.7 ± 0.9; we visually confirmed three sets of *Calptilia* moth species with largely overlapping host use (Fig. 2). The average parasitism rate throughout the year was 46.4%. The parasitism rate for each species is described in Table S3. Six of 13 parasitoid wasps were previously confirmed to have emerged from *Caloptilia* larvae; they provided the reference sequences that were used for constructing *Caloptilia*-blocking primers.

**Figure 2.**
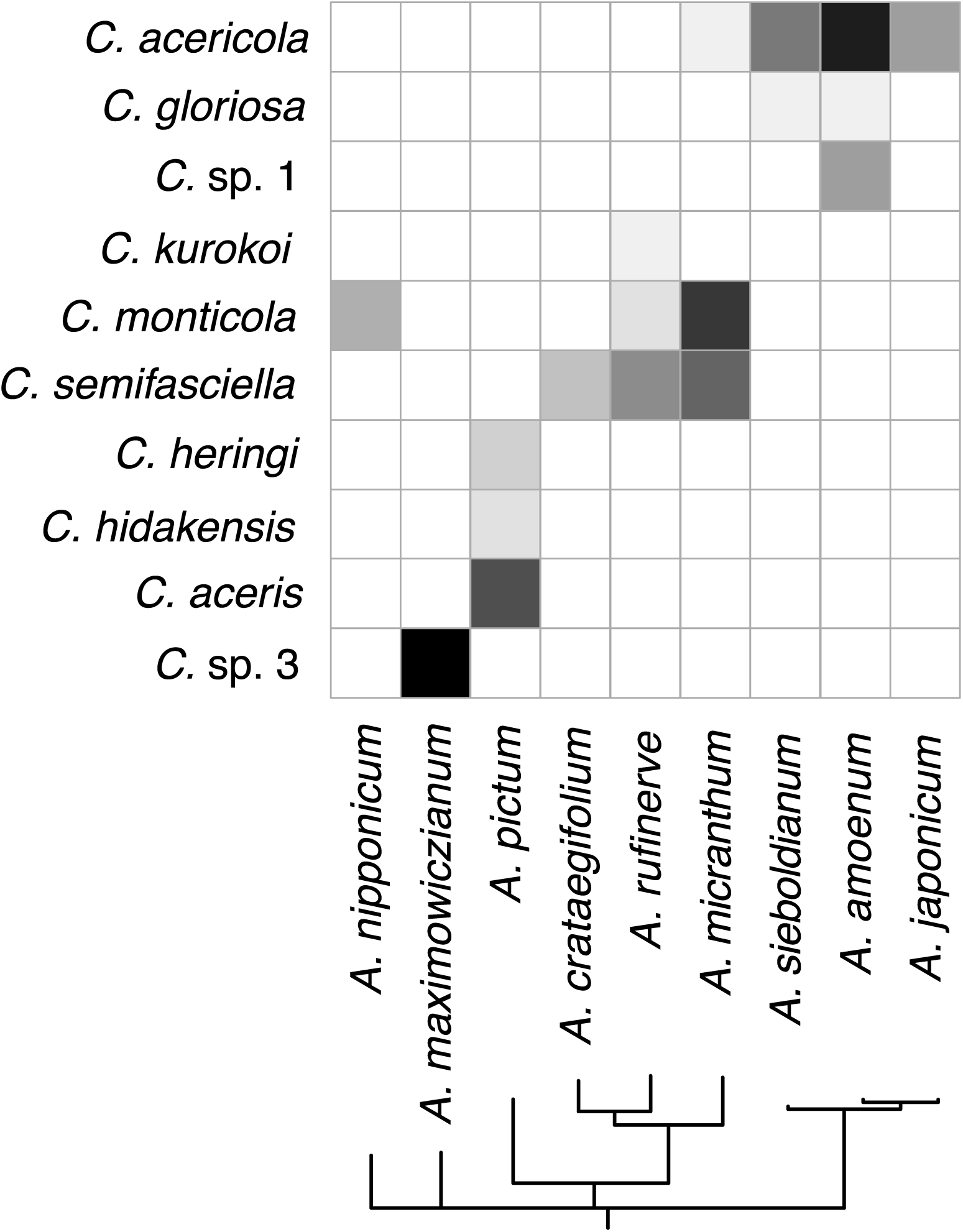
The results of *Acer–Caloptilia* interactions.

**Figure 3.**
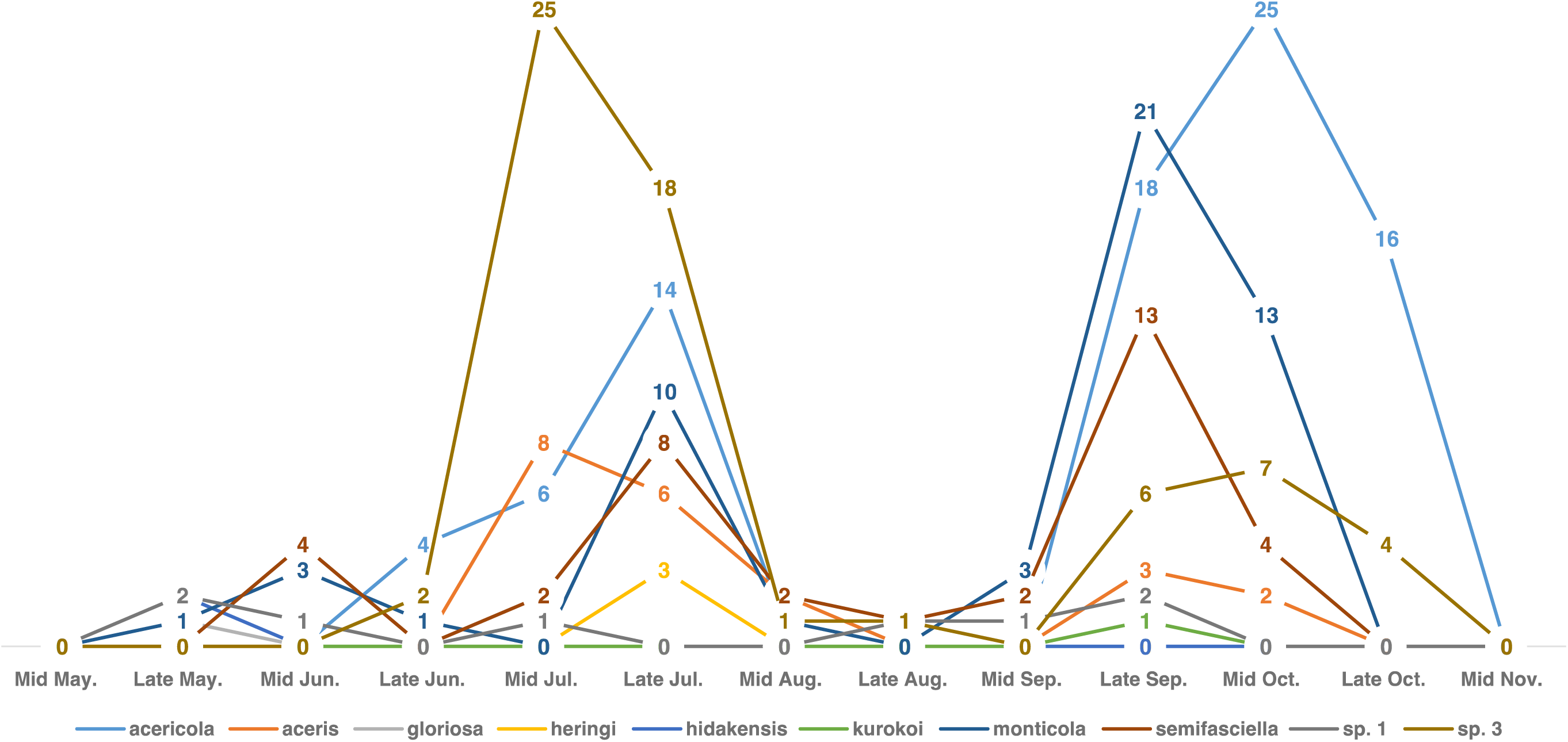
Phenology of 10 *Caloptilia* moths feeding on maples in this study area.

**Figure 4.**
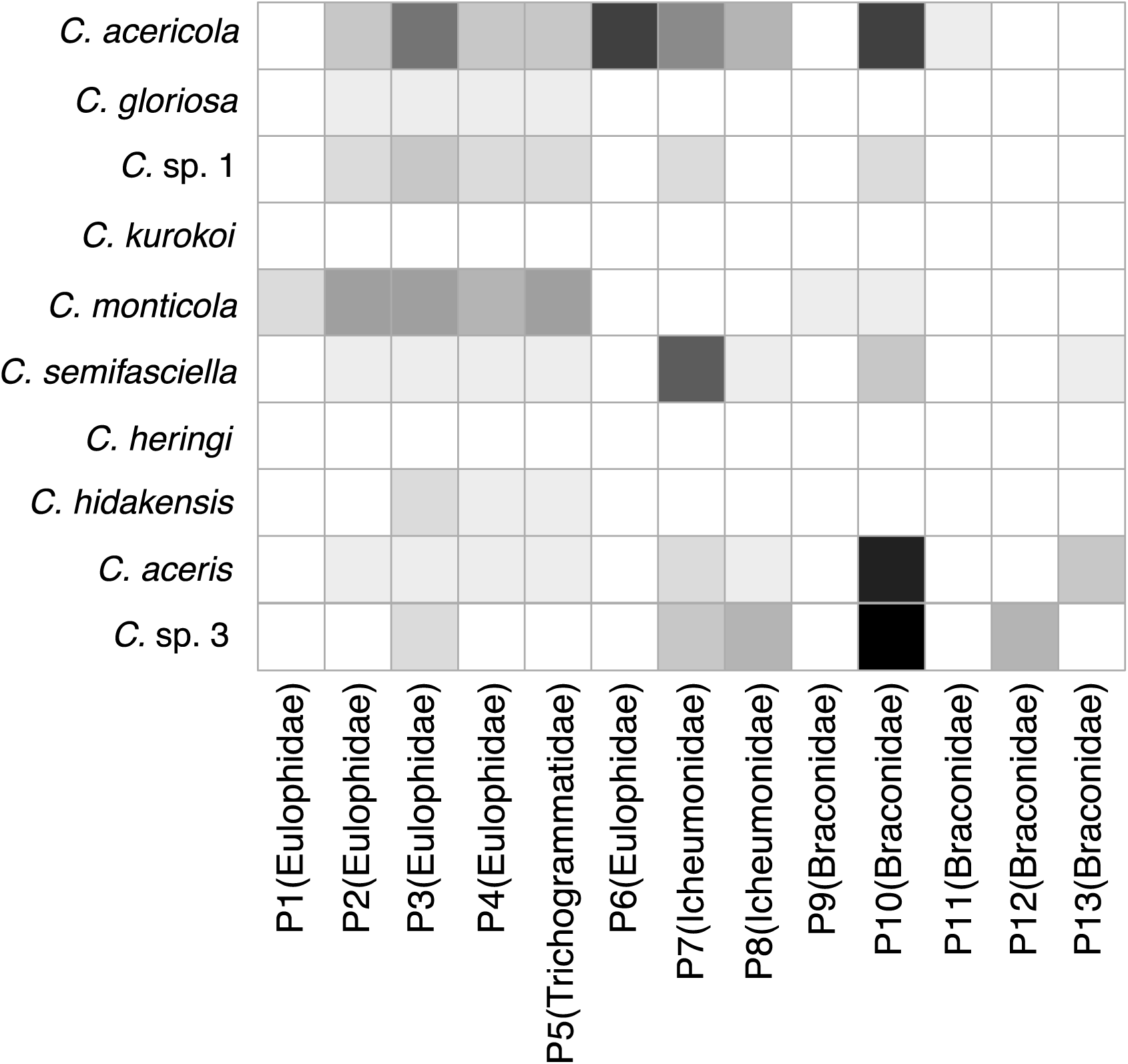
The results of parasitoid wasp–*Caloptilia* moth interactions.

The results of the null model analysis indicated that both temporal and enemy niches showed significantly more overlap among species than the expected random distribution given by both indices (temporal, Pianka *SES* = 4.29, *P* = 0.003, Czechanowski *SES* = 4.77, *P* = 0.001; enemy, Pianka *SES* = 4.77, *P* = 0.001, Czechanowski *SES* = 5.73, *P* = 0.000; Table 1). T This indicates that phenology is significantly overlapping among *Caloptilia* species, and parasitoid wasps are widely shared among *Caloptilia* species. In Mantel tests, only the relationship between temporal and enemy niches, as assessed by the Pianka index, showed a significant correlation (*r* = 0.41, *P* = 0.046; Fig. 5, Table 2), and the Czechanowski index indicated a similar, but not significant, trend (*r* = 0.35, *P* = 0.063; Table 2). Mantel tests showed no significant correlations between other factors (Table 2). This indicates that species with overlapping phenology tend to share common parasitoid wasps.

**Table 1.** The results of comparison with the null model based on 10,000 randomizations. These tests employed Lawlor’s (1980) algorithm RA3, using the R package EcoSimR (Gotelli *et al.* 2013) for enemy niche and the program TimeOverlap, based on the algorithm ROSARIO (Castro-Arellano *et al.* 2010), for temporal niche.

|  | Pianka index |  |  |  | Czechanowski index |  |  |  |
| --- | --- | --- | --- | --- | --- | --- | --- | --- |
|  | Observed | Trend | SES | *P*-values | Observed | Trend | SES | *P*-values |
| Temporal | 0.36 | overlap | 4.29 | **0.003** | 0.79 | overlap | 4.77 | **0.001** |
| Enemy | 0.47 | overlap | 4.77 | **0.001** | 0.39 | overlap | 5.73 | **0.000** |

Bold letters indicate significant results in two-tailed randomization tests

**Figure 5.**
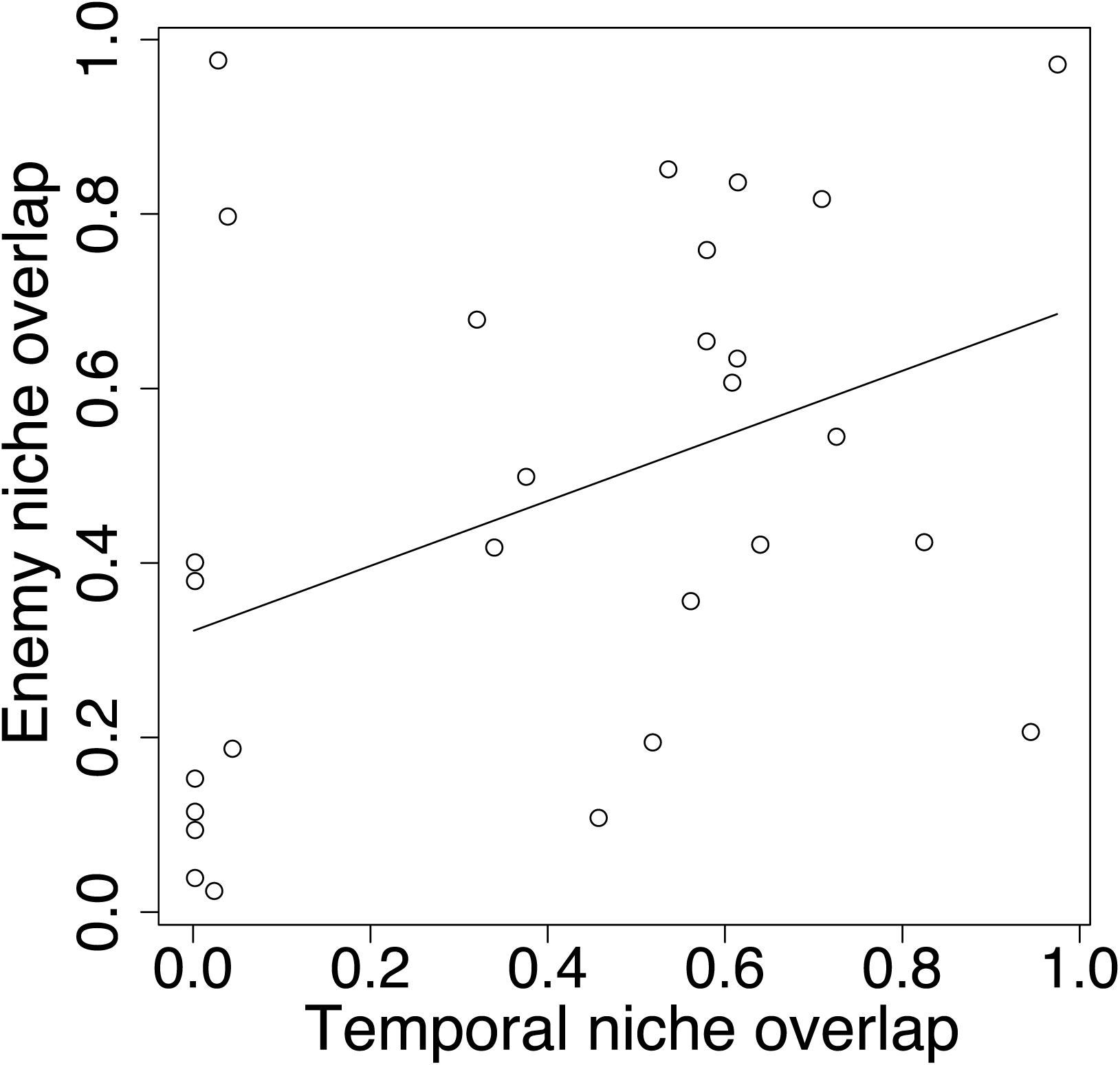
The relationship between the overlaps of temporal and enemy niches in the Pianka index (Mantel *r* = 0.41, *P* = 0.046).

**>Table 2.** Results of Mantel tests of the correlations among resource niche overlap, temporal niche overlap, enemy niche overlap, and phylogenetic distance between *Caloptilia* moths.

|  | Pianka index |  | Czechanowski index |  |
| --- | --- | --- | --- | --- |
|  | Mantel *r* | *P*-values | Mantel *r* | *P*-values |
| Resource-Temporal | 0.00 | 0.973 | -0.06 | 0.690 |
| Resource-Enemy | -0.08 | 0.701 | -0.07 | 0.736 |
| Resource-Phylogeny | 0.14 | 0.244 | 0.19 | 0.154 |
| Temporal-Enemy | 0.41 | **0.046** | 0.35 | 0.063 |
| Temporal-Phylogeny | 0.01 | 0.974 | 0.07 | 0.844 |
| Enemy-Phylogeny | -0.19 | 0.288 | -0.14 | 0.430 |

Bold letters indicate significant results in Mantel tests

## Discussion

### Role of phenology and natural enemies in facilitating species coexistence

The present study found three sets of *Caloptilia* moth species, each consisting of species with largely overlapping host ranges. Although the species that share hosts are not monophyletic, they are very closely related inthe *Caloptilia* phylogeny (except for *C. gloriosa*, which belongs to a different clade than the rest of the maple-feeding *Caloptilia*) and have almost identical larval feeding modes (Fig. S1: Nakadai & Kawakita 2016). Additionally, we found large overlaps in both phenology and parasitoid community among species sharing the same host and among the community of maple-feeding *Caloptilia* as a whole (Table 1). These findings suggest that niche partitioning might not be necessary for closely related herbivores to coexist on shared hosts.

An obvious shortcoming of the above conclusion is that factors not accounted for in our analysis may be critical for niche partitioning among *Caloptilia* species. For example, although there is no apparent difference in the age of leaves used by the larvae or larval feeding mode among the species studied (Fig. 1), there may be a fine-scale difference that we did not detect. Also, because we used larvae at the leaf-rolling stage for our analysis of internal parasitoids, the role of parasitoids at the egg or leaf-mining stage was left uninvestigated. Condon *et al.* (2014) showed that parasitoids often attack the larvae of unusual hosts but do not successfully emerge as adults in such occasions. Because we only searched for parasitoids using the larvae of prey herbivores, such lethal interactions may have been included in the data, obscuring differences in parasitoid communities. Examining every aspect of *Caloptilia* life history may thus reveal an unexpected mechanism that facilitates coexistence of species with overlapping host use.

Alternatively, niche partitioning may genuinely be absent, and species coexistence may be facilitated by other mechanisms. For example, shared natural enemies enhance species coexistence, either if random predation eases interspecific competition among herbivores (Strong *et al.* 1982) or if negative frequency-dependent predation decreases the population of the more abundant species (Ishii & Shimada 2012). Strong (1982) found resource partitioning to be virtually absent among hispine beetles (Chrysomelidae), which commonly coexist as adults in the rolled leaves of *Heliconia* plants, suggesting that pressure from predators and parasites has a stronger influence on community structure in this species than does interspecific competition.

Additionally, Ishii and Shimada (2012) showed that frequency-dependent predation by the pteromalid wasp *Anisopteromalus calandrae* enhanced the coexistence of two bruchid beetles, *Callosobruchus chinensis* and *C. maculatus*. Exploring how parasitoids and other predators (e.g., birds, bats, spiders) control the dynamics of *Caloptilia* populations will be useful in determining whether closely related herbivores can coexist without niche partitioning.

Data on phenological partitioning among closely related herbivores is still sparse (e.g., Yamamoto & Sota 2009), so the generality of temporal niche overlap as observed among the *Caloptilia* species is still unknown. The seasonal dynamics of their food source (maple leaves) may be a straightforward explanation for the observed synchronization of *Caloptilia* phenology, although other abiotic factors, such as temperature or precipitation, may be responsible. In any case, our results strongly indicate that partitioning of phenology is unlikely to be important in facilitating the coexistence of closely related herbivores on shared host plants.

### Assessment of the enemy niche using matabarcoding

Recently, metabarcoding has increasingly been used as a tool for discerning less-visible patterns in food webs (Pompanon *et al.* 2012; Andrew *et al.* 2013; Kartzinel *et al.* 2015; Leray *et al.* 2015). Most studies that employ metabarcoding investigate diet using stomach contents (Kartzinel *et al.* 2015; Leray *et al.* 2015), but such an approach has rarely been used to evaluate parasitoid–prey interactions. Internal parasitoids are usually searched by developing specific primers for each parasitoid taxon (Rougerie *et al.* 2011; Condon *et al.* 2014; Wirta *et al.* 2014). However, metabarcoding allows detection of parasitoid taxa not targeted by specific primers. This approach is particularly useful when the parasitoid community includes taxonomically diverse or unknown species, or when analyzing a large number of prey samples, as in the present study, which involved 274 *Caloptilia* larvae. We also pioneered the use of the blocking primer approach for multiple closely related herbivorous insects by constructing a blocking primer that targets the region in which the sequences are shared only within *Caloptilia*. This method allows metabarcoding to be employed in cases where it is difficult to distinguish between closely related species based on morphology alone. As discussed above, assessment of paraitoid communities solely based on barcoding of herbivore larvae potentially overestimates the breadth of the enemy niche because lethal parasitoid–prey interactions are not omitted from the results. Thus, a combined approach incorporating both barcoding and laboratory rearing will allow a more precise assessment of the enemy niche.

### The link between local species coexistence and global species diversity

Recently, ecologists and evolutionary biologists have recognized that local species coexistence (e.g., current ecological processes) has a major effect on global species diversity (e.g., macroevolutionary outcome) (Rabosky 2009; Tobias *et al.* 2013; Storch *et al.* 2012; Germain *et al.* 2016; Prinzing *et al.* 2016). For example, Prinzing *et al.* (2016) found that angiosperm clades with a greater extent of local co-occurrence are more species rich. To clarify these findings, it is necessary to document examples in other organisms, including herbivorous insects. The results of the present study show that the number of locally co-occurring *Acer*-feeding *Caloptilia* species is a function of both host plant diversity and abundance of species coexisting on the same hosts. Although coexistence of the 10 *Caloptilia* species found in this study is not yet fully explained, improved knowledge of the mechanisms that enable such coexistence is ultimately necessary to our understanding of the processes that generate diversity in herbivorous insects.

## Acknowledgements

We thank staff members of Ashiu Forest Research Station for field assistance, and S. Fujinaga, S. Matsuoka, Y Okazaki and M. Ushio for advicing about DNA experiments. This work was supported by a grant from Grant-in-Aid for JSPS Fellows Grant Number 15J00601 and the JSPS KAKENHI Grant Number 26650165, 15H04421, 15H05241.

## Data Accessibility

Obtained DNA sequences have been deposited in the DDBJ database under accession numbers DRX072707-DRX073080 (BioProject: PRJDB5369) and LC201483–LC201501.

## Author Contributions

R.N and A.K conceived the study and wrote the manuscript.

R.N. designed the study and performed sampling, laboratory work, and data analysis.

## Supporting information

Table S1 OTUs used in the analysis.

Table S2 DDBJ accession numbers.

Table S3 Abundance and parasitoid rate for each species.

Table S4 Information on parasitoid wasps that were used for constructing the blocking primer for *Caloptilia* moths.

Figure S1 Phylogeny of *Caloptilia* moths and their related groups. The phylogeny was constructed by the maximum-likelihood method using four genomic regions (COI, ArgK, CAD, and EF-1a) of 71 species (Nakadai & Kawakita 2016). Red indicates the lineage of *Caloptilia* moths associated with maples.

Supplementary file 1 Fasta-formatted representative sequences of each OTU, created by the UPARSE pipeline.

Supplementary file 2 Performance assessment of the blocking primer created in this study for *Caloptilia* moths associated with maples.

